# Crosslinker-free *in situ* hydrogel induces self-aggregation of human dental pulp stem cells with enhanced antibacterial activity

**DOI:** 10.1101/2024.12.30.630819

**Authors:** Sang Jin Lee, Zhenzhen Wu, Mengyu Huang, Chao Liang, Ziqi Huang, Chen Siyuan, Vidhyashree Rajasekar, Mohamed Mahmoud Abdalla, Haram Nah, Dong Nyoung Heo, Il Keun Kwon, Min-Jai Cho, Seong Jun Kim, Seil Sohn, Su-Hwan Kim, Ryohichi Sugimura, Cynthia Kar Yung Yiu

**Affiliations:** Biofunctional Materials, Division of Applied Oral Sciences and Community Dental Care, Faculty of Dentistry, The University of Hong Kong, 34 Hospital Road, Sai Ying Pun, Hong Kong SAR, PRC; Paediatric Dentistry, Faculty of Dentistry, Prince Philip Dental Hospital, The University of Hong Kong, 34 Hospital Road, Sai Ying Pun, Hong Kong SAR, PRC; Department of Dental Materials, School of Dentistry, Kyung Hee University, 26 Kyungheedae-Ro, Dongdaemun-Gu, Seoul 02447, Republic of Korea; Department of Neurosurgery, Chungbuk National University College of Medicine, Chungbuk National University Hospital, Seowon-gu, Cheongju-si 28644, Chungcheong-do, Republic of Korea; Department of Neurosurgery, CHA Bundang Medical Center, CHA University, Seongnam-si, Gyeonggi-do, Republic of Korea; Department of Chemical Engineering (BK21 FOUR), Dong-A University, Busan 49315, Republic of Korea; School of Biomedical Sciences, Li Ka Shing Faculty of Medicine, The University of Hong Kong, Pokfulam, Hong Kong SAR, PRC; Centre for Translational Stem Cell Biology, Hong Kong SAR, PRC

**Keywords:** *In situ* hydrogel, 3D cell aggregation, antibacterial activity, biocompatible, spontaneous degradation

## Abstract

Recently, injectable hydrogels have garnered significant attention in tissue engineering due to their controlled flowability, strong plasticity, adaptability, and good biocompatibility. However, research on readily injectable *in situ*-forming hydrogels capable of forming functional three-dimensional (3D) tissue condensations remains limited. This study explores the development and evaluation of a carboxymethyl chitosan (CMCTS) / oxidized hyaluronic acid (oHA) hydrogel incorporated with silver sulfadiazine (AgSD) for tissue engineering applications with inherent antibacterial activity. Through physicochemical analysis, the optimal formulation of CMCTS/oHA hydrogels was established. The hydrogel demonstrated excellent injectability, enabling minimally invasive *in situ* delivery. *In vitro* cytotoxicity assays identified 0.1% AgSD as the optimal concentration, supporting cell proliferation while exhibiting antimicrobial efficacy against *S. mutans* and *E. faecalis*. *In vivo* studies revealed complete hydrogel degradation and good biocompatibility, with no adverse tissue reactions. The hydrogel’s ability to form 3D cell aggregates and support tissue regeneration underscores its potential for future 3D tissue engineering applications. Consequently, the injectable CMCTS/oHA/AgSD hydrogel developed in this study holds significant potential for application in a wide range of bioengineering fields, including antibacterial substance delivery systems and 3D tissue engineering, indicating potential for future clinical application.

## 1. INTRODUCTION

Human body tissues can be unexpectedly damaged by severe trauma, injury, and tumor resection [1]. Over the past decade, various novel tissue engineering approaches have aimed to restore damaged tissues by leveraging biomaterials, cells, and growth factors to stimulate repair and regeneration [2]. Key strategies include scaffold-based tissue replacement, cell-based therapies, and the use of bioactive molecules to enhance the body’s natural healing processes [3].

Among the biomaterial approaches for recovering damaged tissue, injectable hydrogel systems have become a widely used scaffold approach [4]. A key advantage is their ability to be delivered in a minimally invasive manner, allowing the hydrogel to conform to complex tissue geometries at the injury site [5]. Injectable hydrogels can also be engineered to incorporate bioactive factors, such as growth factors or therapeutic drugs, in addition to stem cells, to stimulate and guide the tissue regeneration process [6]. Various types of injectable hydrogels have been investigated, including supramolecular peptide hydrogels [7] and thermos-responsive hydrogels [8]. Although these hydrogel systems have shown excellent performance in terms of regenerative outcomes, they face several challenges. Peptide-based hydrogels, for example, can be expensive to prepare, resulting in limitations with mass production. Thermo-responsive hydrogels form in response to temperature changes, but this process can take several minutes for solid gelation, which can result in discomfort for patients waiting in the surgery table or treatment chair. This requires a holding period, and the hydrogel usually forms from an aqueous solution when the body temperature is maintained at an appropriate level, meaning they cannot form hydrogels at room temperature. In the case of ultraviolet (UV)-based hydrogel, it starts as a soluble hydrogel and requires UV crosslinking after being applied to the target area, making it challenging to use *in situ* as immediate injectable form [9]. These deficiencies hinder the convenience of loading various molecules or transplanting stem cells within the hydrogel for rapid delivery, leading to significant limitations in clinical applications for swift treatment.

As a regenerative medicine’s tool, stem cell therapy also plays a crucial role in inducing the regeneration of damaged tissues [10]. In particular, the implantation of 3D tissue constructs (e.g., 3D spheroids or aggregates) has emerged as a state-of-the-art system in the field of tissue engineering and regenerative medicine [11, 12]. These 3D tissue-engineered grafts offer significant advantages over traditional cell-based therapies, as they can more closely mimic the native extracellular matrix and cellular interactions found in complex tissues [13]. 3D tissue platforms are playing a transformative role in advancing regenerative strategies across a range of clinical applications [11]. Many researchers have investigated the use of hydrogels to pursue 3D tissue remodeling through cell condensation-laden hydrogel systems [14, 15]. For instance, Zhang *et al.* developed extracellular matrix hydrogels and manually embedded pre-made 3D spheroids, which were then implanted subcutaneously in mice [16]. In another study, Kwon *et al.* fabricated spheroid-laden hydrogels, showing potential *in vivo* applications in tissue engineering and regenerative medicine [17]. While the transplantation of 3D stem cell aggregates has demonstrated promising regenerative potential, these systems still encounter certain drawbacks. Notably, the fabrication of these constructs requires a cumbersome cross-linking step using complex apparatus prior to implantation [18]. Additionally, the residual hydrogel remaining after transplantation can impede the integration of the graft with the host tissue [19]. To eliminate the long-term degradation of these persistent hydrogel materials, the use of enzymes or other degradative agents is necessary, which requires an additional inconvenient step [20]. These drawbacks associated with the preparation and post-implantation behavior of hydrogel-based stem cell constructs present areas for further optimization and innovation.[21]

Therefore, developing more user-friendly and conveniently manageable hydrogel fabrication methods that allow for smoother integration with the host tissue could help overcome these challenges and unlock the full regenerative capabilities of advanced 3D tissue engineering approaches. Unfortunately, a convenient injectable system that can organize stem cells into tissue-like constructs and allow for the natural biodegradation of the hydrogel has not yet been clearly established.

To address these issues, we have designed a crosslinker-free and high-density dental pulp stem cells (DPSCs)-laden *in situ* injectable hydrogel system, where the hydrogel is formed through the self-assembly of a macromer solution at room temperature, enabling the spontaneous 3D organization of stem cells during hydrogel degradation. In our previous reports, we successfully prepared a self-crosslinkable hydrogel using water-soluble glycol chitosan (gC) and oxidized hyaluronic acid (oHA) at room temperature [22, 23]. This hydrogel system has demonstrated promising *in vitro* and *in vivo* results when used for *in situ* therapeutic applications, as evidenced by its implantation in a calvarial defect model [22] and a spinal cord injury model [23] in rats.

Building on these prior findings, in this study, we propose the use of high-density DPSCs encapsulated within a carboxymethyl chitosan (CMCTS) and oHA hydrogel system, augmented with antibacterial silver sulfadiazine (AgSD) materials. As cell source, this study selected DPSCs because they are commonly used in craniofacial stem cell research, garnering significant attention in regenerative medicine due to their unique properties. These properties include their ability to self-renew, multilineage differentiation potential, and immunomodulatory capabilities [24]. Additionally, DPSCs demonstrate strong cell-to-cell communication, which promotes their self-aggregation [25]. Given these regenerative characteristics, DPSCs have been effectively integrated with various biomaterials to develop biomimetic constructs for cell therapy applications [26] and obtained from discarded wisdom teeth, allowing us to extract valuable cells from waste resources [27]. The AgSD was incorporated into the hydrogel crosslinking process to demonstrate the substance delivery capabilities of hydrogels and to enhance the antimicrobial properties of the constructs [28, 29]. Our detailed strategy is illustrated in Fig. 1. The CMCTS, oHA, DPSCs, and AgSD were directly conjugated to form the hydrogel construct at room temperature (Fig. 1a). Next, 3D cell formulation, cell viability, and antibacterial activity were characterized (Fig. 1b), followed by the direct injection of the optimal CMCTS/oHA/AgSD hydrogels into mice subcutaneously without incision to verify self-degradation and cytocompatibility (Fig. 1c).

**Fig. 1.**
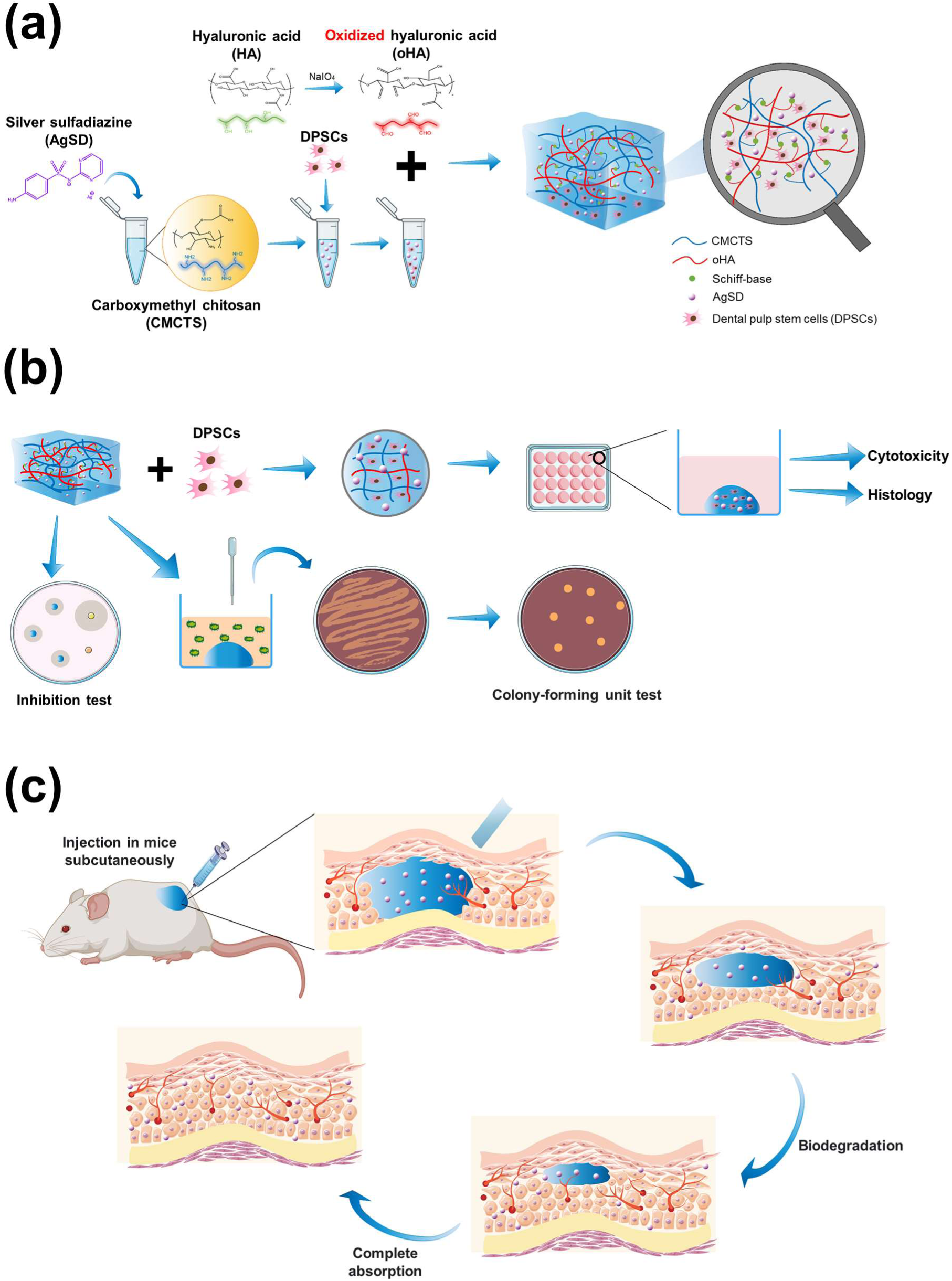
Schematic illustration of the experimental setup for AgSD-incorporated CMCTS/oHA injectable hydrogel and its *in vitro* and *in vivo* tests. AgSD was incorporated into the CMCTS macromer solution, followed by conjugation with oHA to fabricate CMCTS/oHA/AgSD hydrogels (a). High-density DPSCs-laden CMCTS/oHA/AgSD hydrogels were cultured for 3D cell condensation, and the CMCTS/oHA/AgSD hydrogels were analyzed using inhibition zone and colony-forming unit tests to assess antibacterial activity (b). Injectable CMCTS/oHA/AgSD hydrogels were injected subcutaneously into mice to evaluate degradation and biocompatibility (c).

## 2. Materials & methods

### 2.1 Materials

O-carboxymethylated chitosan (CMCTS, Cat. No. sc-358091) with a deacetylation degree of ≥ 90% was purchased from Santa Cruz Biotechnology (Santa Cruz, CA, USA). Silver(I) sulfadiazine (AgSD, Cat. No. 481181), ethylene glycol (Cat. No. 102466), and deuterium oxide (D2O, Cat. No. 151882, isotopic purity of 99.9 atom% D) were purchased from Sigma-Aldrich (St. Louis, MO, USA). High-molecular-weight hyaluronic acid powder (HA, 1000 kDa, Cosmetic & Food Grade) was purchased from The Pharmers Market (Manchester, United Kingdom). A Luer-lock tip 1.0-mL syringe (Cat. No. 309628) was purchased from BD Biosciences (Becton Dickinson, USA). A tunnel mixer (Cat. No. MT-SEM01) to connect two syringes was purchased from NOBAMEDI Co., Ltd. (Yongin, Republic of Korea).

### 2.2 Preparation of CMCTS/oHA and CMCTS/oHA/AgSD hydrogels

#### 2.2.1 Preparation of oHA with sodium periodate (NaIO_4_)

According to our previous literature [22, 23], oHA was prepared with certain modifications using NaIO_4_ in the oxidation process. Briefly, 3.8 g of HA was added to 360 mL of deionized water and stirred until completely dissolved at room temperature. Afterward, 1.068 g of NaIO_4_, pre-dissolved in 40 mL of Milli-Q water, was added to the HA solution with vigorous stirring in a dark place overnight. Next, an excess quantity of ethylene glycol (1 mL) was added to the solution to terminate the reaction for 30 minutes at room temperature. To eliminate any remaining compounds, the solution was dialyzed for 7 days using a dialysis bag (Cat. No. JHC0095, 8000-14000 MWCO, BioMomei, China) with deionized water replacement every 12 hours. Finally, the purified solution was lyophilized for 1 week to obtain the clear oHA composite. The lyophilized composite was kept at -20 °C until use.

#### 2.2.2 Preparation of CMCTS/oHA and CMCTS/oHA/AgSD hydrogels

To optimize the hydrogel fabrication method, CMCTS/oHA was fabricated using two different preparation methods (tunnel mixer or pipette mixing) and two different concentrations of CMCTS. Briefly, 2% and 4% CMCTS composites were dissolved in PBS (weight/volume), and 3% oHA composite was dissolved in PBS (weight/volume), separately. After fully dissolving by vortexing, the 2% or 4% CMCTS and 3% oHA macromer solutions were uniformly mixed by tunnel mixer or pipette mixing methods on clean petri dishes.

To determine the optimal hydrogel ratio, we prepared hydrogels of CMCTS/oHA in ratios of 2:8, 5:5, and 8:2, and measured the gelation time and formation probabilities. First, we prepared 4 wt% of CMCTS and 3 wt% of oHA. Different ratios of the hydrogel components were created and mixed using a hand-mixing method, with a total volume of 200 μL for each condition (n=4). Gelation time was measured by observing when the CMCTS/oHA hydrogels were fully formed, with no residual macromer solution remaining on the dish. After the hydrogels formed, all samples were lyophilized using a freezing dryer for 3 days. The cross-sectional morphology was characterized by scanning electron microscopy (SEM, SU1510, Hitachi, Japan). Based on these measurements, the CMCTS/oHA ratio of 8:2 with tunnel mixing was chosen for further *in vitro* and *in vivo* study. For the preparation of AgSD-loaded CMCTS/oHA hydrogels, a series of AgSD (0.1, 0.25, 0.5%, wt/v) were added to the CMCTS macromer solution followed by the addition of oHA to produce CMCTS/oHA + 0.1% AgSD, CMCTS/oHA + 0.25% AgSD, and CMCTS/oHA + 0.5% AgSD hydrogels through a tunnel mixer with an 8:2 (CMCTS/oHA) ratio, respectively.

### 2.3 Characterization of CMCTS/oHA hydrogels

#### 2.3.1 ^1^H-Nuclear magnetic resonance and fourier-transform infrared spectroscopy

To confirm the successful synthesis of oHA, a 1H-Nuclear Magnetic Resonance (NMR) spectrometer (Bruker Avance 400, Bruker Corporation, Billerica, MA, USA) was used to measure the chemical structure of oHA with D2O as the solvent. Furthermore, the characteristic peaks in the infrared spectrum of CMCTS and oHA composites, and lyophilized CMCTS/oHA composite from hydrogel samples were measured by Fourier-Transform Infrared (FT-IR) spectrometer (Spectrum Two™, PerkinElmer, Norwalk, CT, USA) with the wavenumber ranging from 500 to 4000 cm_−1_.

#### 2.3.2 Rheology

The hydrogel-forming kinetics and mechanical properties of CMCTS/oHA hydrogels were tested using an MCR 92 Rheometer (Anton Paar, GmbH, Germany) equipped with parallel plates in oscillatory mode at room temperature and 37 °C. Briefly, two different concentrations of CMCTS solution (2% and 4%) were mixed with 3% oHA solution by two different methods (tunnel mixer and pipette mixing). Subsequently, hydrogels were measured using a rotating rheometer with a plate-plate geometry of 8 mm diameter and a 1 mm gap to investigate the frequency-dependent viscoelastic behavior. Under controlled conditions of room temperature and steady shear flow procedure, the storage elastic modulus (G′) and loss modulus (G″) values of each sample were measured. The frequency sweep was carried out in the range of 0.1 - 10 Hz at a constant strain of 1%.

#### 2.3.3 *In vitro* degradation of lyophilized CMCTS/oHA composites

The 2% CMCTS/3% oHA hydrogels and 4% CMCTS/3% oHA hydrogels were prepared and weighed, recording the initial weight (W₀) after 48 hours of lyophilization. The freeze-dried samples were placed in 12-well culture plates (Cat. No. CS016-0093, ExCell Bio, Shanghai, China) and immersed in 3 mL phosphate-buffered saline (PBS) solution (pH = 7.4). The culture plates were placed in a 5% CO_2_ incubator at 37 °C and the lyophilized hydrogels were shaken in an orbital incubator at 80 rpm (Cat. No. Stuart SI50, Rhys International Ltd., Bolton, UK). The PBS in the plates was carefully replaced every day. At predetermined time points (1, 2, 3, 5, 7, and 14 days, n = 4 for each timepoint), the hydrogel samples were collected and washed twice with deionized water before freeze-drying for 48 hours. The weight measurements and photographs were taken to record changes in mass and shape over time. The weight after degradation at specific intervals (W_t_) was used to calculate the (1) degradation rate and (2) residual mass of these two types of hydrogels using the following formulas:

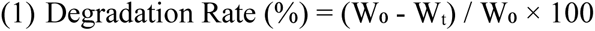

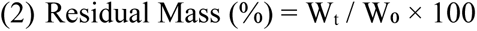

Additionally, to determine changes in the internal microstructure of hydrogels during the degradation process, the hydrogel with better properties (4% CMCTS / 3% oHA) was examined by scanning electron microscopy (SEM, Cat. No. SU-1510, Hitachi High-Technologies, Tokyo, Japan). Specifically, hydrogels immersed in PBS at different time intervals were collected, lyophilized after liquid nitrogen pre-freezing, and cut longitudinally to allow cross-sections. A thin Au film was sputter-coated onto the cross-sections, and the change in interior microstructure was observed by SEM characterization. The average pore size of hydrogels at different time points was calculated and analyzed using ImageJ software based on SEM images.

#### 2.3.4 *In vitro* degradation of CMCTS/oHA hydrogels under different medium conditions

We further performed the degradation test under several different kinds of conditions such as 1) 1x PBS, 2) 10% FBS, and 3) completed cell growth medium. Briefly, 4% CMCTS/ 3% oHA hydrogels were placed into a 12-well plate directly and 3 ml of mediums were added to immerse the hydrogel completely (n=3 for each group). After that, the plate was located into the incubator (TT-60-SI InQShake Shaking Incubator, Hercuvan lab systems, United Kingdom) and shaken for 10 days under 5% CO_2_, 37 °C and 80 rpm. In order to maintain a constant concentration, it is essential to replace the solution and take images daily. Eventually, the volumes of hydrogels in different mediums were calculated using polygon selections and measurement functions in ImageJ software based on the hydrogel outlines from daily images. The volume changes over time for different groups were analyzed using Prism 10 software.

#### 2.3.5 Morphology

Based on the rheology and degradation results in Sections 2.3.2 and 2.3.3, 4% CMCTS/3% oHA hydrogels were chosen for subsequent experiments. To determine whether the addition of AgSD affects the morphological characteristics of the hydrogel, CMCTS/oHA and CMCTS/oHA + 0.5% AgSD hydrogels were prepared under the same conditions and imaged to observe their morphologies. The hydrogels were then frozen at -80 °C overnight, followed by lyophilization for 48 hours. The surface and interface characteristics of these hydrogels were assessed by photographs and SEM.

### 2.4 *In vitro* cell test

The DPSCs were purchased from Lonza (Cat. No. PT-5025, Basel, Switzerland) and cultured using alpha modified minimal essential medium (α-MEM, Cat. No. 11900-024, Thermo Fisher Scientific Inc., Waltham, MA, USA) supplemented with 10% fetal bovine serum (FBS, Cat. No. 10270106, Thermo Fisher Scientific Inc.) and 1% penicillin-streptomycin (P/S, Cat. No. 15140163, Thermo Fisher Scientific Inc.) at 37 °C with 5% CO_2_. Passages from two to four were used in experiments. After the cells were expanded in T-75 flasks, they were trypsinized using 0.25% trypsin-EDTA (Cat. No. 25200072, Thermo Fisher Scientific Inc.). The trypsinized DPSCs pellets were directly encapsulated in the CMCTS or CMCTS/AgSD macromer solutions at a density of 100 x 10⁶ cells/mL of total hydrogel before conjugation with oHA. The detailed hydrogel preparation process is described in Section 2.2.

### 2.5 Antibacterial activities of CMCTS/oHA/AgSD

#### 2.5.1 Zone of inhibition test

The antibacterial activity of the CMCTS/oHA/AgSD hydrogel was tested using *Streptococcus mutans* (*S. mutans*) ATCC 700610 and *Enterococcus faecalis* (*E. faecalis*) ATCC 29212 strains, which are commonly found in dental caries and root canals. Samples of CMCTS/oHA/AgSD hydrogel with varying concentrations of AgSD were irradiated with UV light for 30 minutes in an ultra-clean workbench before use. For the assay, 50 μL of each bacterial suspension was inoculated onto Mueller-Hinton agar (Cat. No. CM0337, Oxoid™, Basingstoke, Hampshire, UK) medium at a concentration of 1 × 10^6^ colony-forming units (CFU) / mL. Five uniform circular wells were created in each agar plate using 200 μL pipette tips. Subsequently, 50 μL of five different samples were added into the wells. The samples included: 0.5 mg/mL TAP (Triple Antibiotic Paste, combining Ciprofloxacin (Cat. No. 17850, Sigma-Aldrich), Minocycline (Cat. No. M9511, Sigma-Aldrich), and Metronidazole (Cat. No. M1547, Sigma-Aldrich) in a 1:1:1 ratio) as the positive control; PBS as the negative control; CMCTS/oHA hydrogel; CMCTS/oHA + 0.1% AgSD hydrogel; and CMCTS/oHA + 0.25% AgSD hydrogel. The agar plates were then sealed and incubated overnight in an anaerobic chamber (Baker Ruskinn, Concept Ruskinn, Bridget, UK). The diameter of the bacterial inhibition zones was measured and photographed. The experiments were independently repeated three times to ensure reliability.

#### 2.5.2 CFU test

The antibacterial activity of the CMCTS/oHA/AgSD hydrogel was further evaluated using the CFU test. For this test, 150 μL of 10^6^ CFU/mL *S. mutans* and *E. faecalis* bacterial suspensions were co-cultured with 100 μL of each sample for 24 hours. The samples included: 0.5 mg/mL TAP (Triple Antibiotic Paste, combining Ciprofloxacin, Minocycline, and Metronidazole in a 1:1:1 ratio) as the positive control; PBS as the negative control; CMCTS/oHA hydrogel; CMCTS/oHA + 0.1% AgSD hydrogel; and CMCTS/oHA + 0.25% AgSD hydrogel. Following co-culturing, the bacterial suspensions were serially diluted 10,000 times with Bacto Brain Heart Infusion (BHI, Cat. No. CM1135, Oxoid™) and inoculated onto solid agar culture medium dishes (Cat. No. CM0337, Oxoid™). The agar plates were then sealed and incubated in an anaerobic chamber for 2 days to allow for bacterial colony formation. The number of bacterial colonies was subsequently counted to evaluate the antimicrobial effect of the different samples. The experiments were independently repeated three times to ensure reliability.

### 2.6 *In vivo* test

All animal studies were conducted under the animal research protocol (No. 22-268) approved by the Committee of the Use of Live Animals in Teaching and Research (CULATR) at the University of Hong Kong. The studies adhered to the Animals (Control of Experiments) Ordinance (Hong Kong) and guidelines from the Centre for Comparative Medical Research (CCMR), Li Ka Shing Faculty of Medicine, The University of Hong Kong. Male BALB/c mice (6-8 weeks old) were used for the CMCTS/oHA hydrogel experiments. The mice were obtained from the Experimental Animal Center of the University, with access to food and water ad libitum, and were maintained under pathogen-free conditions. All procedures were performed under anesthesia using intraperitoneal injections of ketamine/xylazine mixtures (100 mg/kg ketamine (Cat. No. 013004, AlfaMedic Ltd.) + 10 mg/kg xylazine (Cat. No. 013006, AlfaMedic Ltd.). Anesthesia was confirmed by checking body reflexes, and eye ointment (Duratears® ointment, Cat. No. 05686, Alcon, Fort Worth, TX, USA) was applied to prevent blindness. The subcutaneous injection sites on the mice were decontaminated with alternating applications of 70% alcohol and povidone-iodine (Betadine®) using sterile cotton swabs. CMCTS/oHA/AgSD hydrogels were then injected into the left and right backs of the mice, with 500 μL volumes per site. The mice were placed in an intensive care unit (ICU) until they fully recovered, after which they were returned to their original housing location. Seven days post-injection, the mice were euthanized by an overdose of Pentobarbital (250 mg/kg) (Dorminal®, Cat. No. 013003, AlfaMedic Ltd.) administered intraperitoneally. The injection areas were harvested and immediately fixed in 10% neutral-buffered formalin (NBF) for 24 hours for histological analysis. The samples underwent a series of ethanol, xylene-ethanol, and xylene treatments for 2 minutes each, then were embedded in paraffin and sectioned at 10 μm thickness using a rotary microtome (Cat. No. RM2155, Leica Microsystems, Wetzlar, Germany). Sections were deparaffinized, rehydrated, and stained with hematoxylin (Cat. No. SH4777, Harris Hematoxylin, Cancer Diagnostic Inc., Durham, NC, USA) and eosin (Cat. No. CS701, Dako, Glostrup, Denmark) (H&E), alcian blue (pH 2.5) with 0.1% nuclear fast red (Cat. No. PH1082, Scientific Phygene®, Fuzhou, China), and 0.5% safranin O (Cat. No. 02782-25, Polysciences Inc., Warrington, PA, USA) with fast green (Cat. No. PH1852, Scientific Phygene®).

### 2.7 Statistical analysis

All values are depicted as mean ± standard deviation. Statistical analysis of the cell proliferation (Fig. 4b and 5b) and antibacterial inhibition zone test (Fig. 6b) were performed using two-way analysis of variance (ANOVA) with Bonferroni *post-hoc* test in GraphPad^®^ Prism 5.0 (GraphPad Software, San Diego, CA). Statistical analysis of the antibacterial CFU test (Fig. 6e and f) and gelation time (Fig. S1b) were performed using one-way analysis of variance Bonferroni *post-hoc* test in GraphPad^®^ Prism 5.0.

## 3. Results and Discussion

The oHA was synthesized by oxidatively cleaving the cyclic structure of hyaluronic acid (HA) into an open chain with sodium periodate, generating two aldehyde groups from the vicinal diol units of the HA main chains (Fig. 2a) [30]. The chemical structure of oHA was confirmed by ^1^H-NMR analysis. As shown in Fig. 2b, the peaks at 4.9 ppm, 5.0 ppm, and 5.1 ppm represent the hemiacetal protons of oHA, which indicates that the aldehyde groups were successfully synthesized through the oxidation process [30, 31]. Consequently, the CMCTS/oHA hydrogel was formed by a Schiff-based reaction between the amino groups of CMCTS and the dialdehyde groups of oHA (Fig. 2c) [32, 33].

**Fig. 2.**
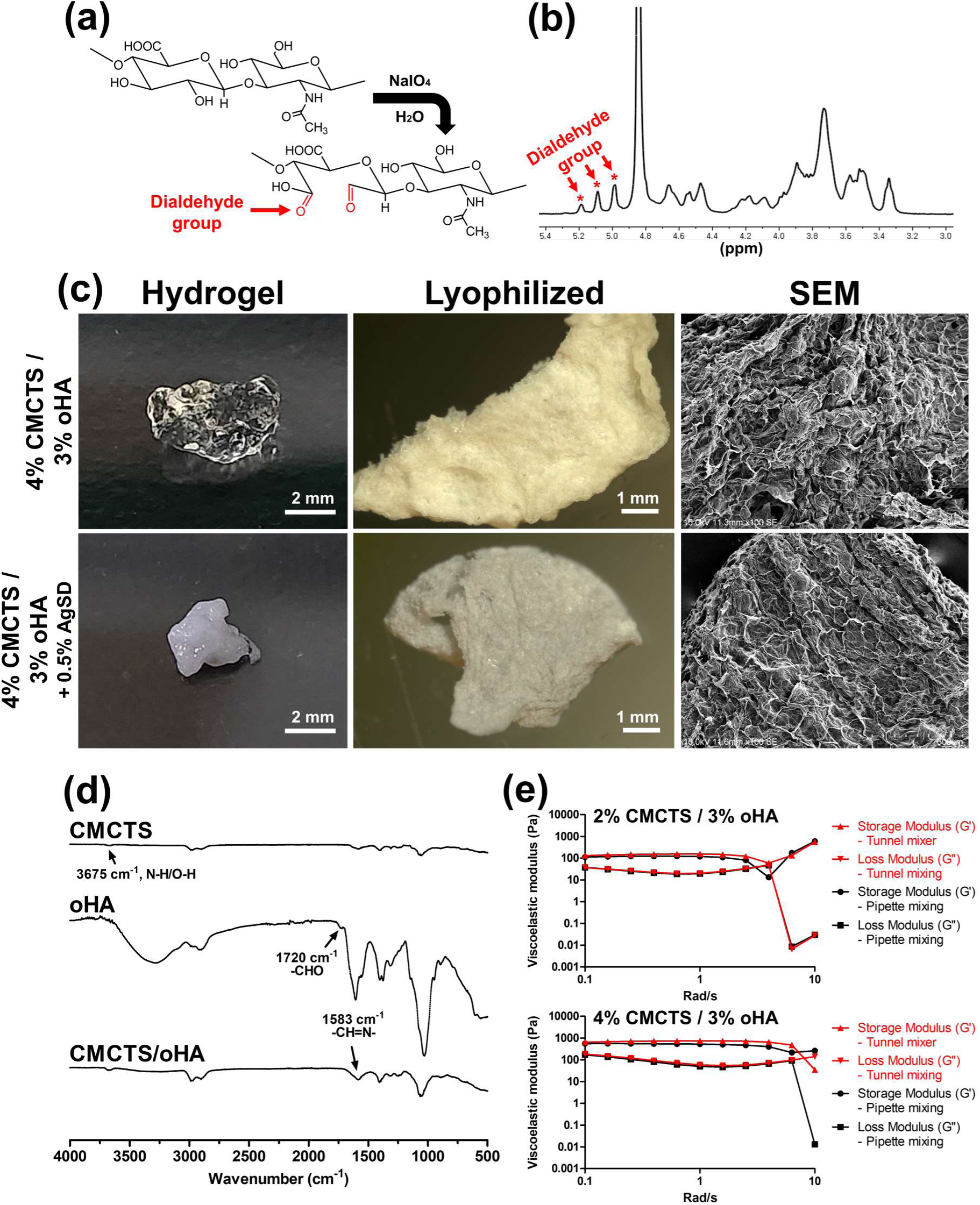
Characterization of CMCTS/oHA and CMCTS/oHA/AgSD hydrogels. Schematic illustration of oHA preparation through NaIO_4_ modification (a). ^1^H-NMR results of oHA indicating the appearance of dialdehyde groups in the oHA macromer solution (b). Fabricated CMCTS/oHA and CMCTS/oHA + 0.5% AgSD hydrogels (c). The hydrogels were well-formed and lyophilized for SEM analysis, exhibiting a uniform porous structure on the surface. FT-IR analysis of original CMCTS powder, synthesized oHA composite, and lyophilized CMCTS/oHA hydrogels (d). Rheology analysis of 2% CMCTS / 3% oHA and 4% CMCTS / 3% oHA hydrogels fabricated via tunnel mixer and pipetting mixing (e). Both methods did not affect the hydrogel properties and showed no differences.

In this study, we optimized the ratio of CMCTS/oHA hydrogel to 8:2 based on our supplementary analysis (Fig. S1). We prepared mixtures of 4% CMCTS and 3% oHA in three different combinations: 2:8, 5:5, and 8:2 (Fig. S1a). The gelation times of these mixtures were presented in Fig. S1b. The results showed that the 2:8 ratio of CMCTS/oHA required almost 2 minutes to completely form the hydrogel. In contrast, the 5:5 and 8:2 ratios achieved gelation in 30 seconds, indicating that these two ratios are preferable for hydrogel formation. Images of hydrogel morphology indicated that the 8:2 ratio was optimal, as there was no residual water around the hydrogels after formation. In comparison, the other two groups exhibited significant residual liquid, suggesting inadequate crosslinking of the polymer chains, which adversely affected their performance [34]. Additionally, cross-sectional SEM images illustrated that the internal pore morphology of the 8:2 ratio is much more regular than that of the other two ratios, with more uniform pore structure (Fig. S1a). Regular and uniform pores facilitate cell aggregation and the formation of 3D tissues in cell-laden hydrogels [35–37]. Based on the results, we determined 8:2 ratio could maximize stem cell encapsulation efficiency due to rapid formulation without residues. The slow hydrogel formation of the 2:8 condition is regarded to the significantly lower presence of amide groups from CMCTS compared to the dialdehyde groups from oHA, which likely unable hydrogel formation. For similar reasons, we utilized a crosslinked hydrogel with a higher glycol chitosan content and lower oHA content in our previous research [22, 23].

The conditioned hydrogel generation process is depicted in Fig. S2. The morphological analysis of the hydrogels was conducted using macroscopic photographs and SEM characterization. We also attempted to add AgSD to the 4% CMCTS/3% oHA, which resulted in a significant color change due to the reaction. Color change fundamentally arises from the reaction between the amino groups (-NH_2_) in CMCTS and the dialdehyde groups (-CHO) in oHA, forming Schiff base bonds and thereby facilitating cross-linking [38]. Silver (Ag) ions can coordinate with functional groups, such as the amino groups in the hydrogel, establishing connections between polymer chains and increasing the cross-linking density of the hydrogel [39]. The AgSD acts as a small molecule catalyst, accelerating the formation of imine bonds. This approach addresses the issues of poor structural stability and weak mechanical properties in the hydrogel [40]. Under conjugation, surface plasmon resonance occurs on the surface of silver nanoparticles (AgNPs), enabling these nanoparticles to absorb and scatter light, resulting in specific colors (Fig. 2c and S2). As the concentration of Ag ions in the hydrogel increases, the number of AgNPs correspondingly rises, leading to greater absorption and scattering of light and resulting in a deeper color of the hydrogel [41]. The existence of AgSD was clearly observed in sectioned staining images (Fig. S3). After lyophilization, the photographs and SEM images showed that the inclusion of AgSD had minimal impact on the surface and interface of the hydrogel, as well as its original porous structure at both the macro and micro levels. Furthermore, the hydrogel formulation was characterized by FT-IR spectrum (Fig. 2d). In contrast to oHA, the characteristic absorption peak for the aldehyde group at 1720 cm^-1^ vanished from the CMCTS/oHA hydrogel, while the imine stretching vibration absorption peak appeared at 1583 cm^-1^ [33], illustrating the formation of dynamic imine bonds between CMCTS and oHA [32].

The sol-to-gel transition of hydrogels was demonstrated by the relative values between *G’* and *G’’*. Fig. 2e showed that both conditions exhibited semisolid hydrogels when the frequency was moderate (less than 6 Hz) because the energy storage modulus *G’* was greater than the loss modulus *G’’*. At this point, the hydrogels displayed the primary elastic characteristics of gels, allowing them to partially withstand external forces. However, *G’’* of the 2% CMCTS/3% oHA condition progressively exceeded *G’* and approached a solution state when the frequency increased to 6 Hz, whereas the elastic behavior of the 4% CMCTS/3% oHA condition remained relatively stable until the frequency approached 10 Hz. Furthermore, the 4% CMCTS/3% oHA hydrogels had much larger elastic and loss moduli than the 2% CMCTS/3% oHA condition, which is related to the number of potential crosslinking sites per chain and the increased internal crosslinking density with a higher concentration of carboxymethyl chitosan [42–44]. In each condition, the two different preparation methods (tunnel and pipette mixing) had little impact on the rheological properties of the hydrogels. Each type of hydrogel had a *G’* value no more than two orders of magnitude larger than *G’’*, indicating that the hydrogel’s phenomenon was weak, providing a great potential for producing injectable hydrogels rather than viscoelastic solid hydrogels from these two systems [45].

The *in vitro* degradation properties of lyophilized 2% CMCTS/3% oHA and 4% CMCTS/3% oHA composites were characterized by macroscopic photographs, residual mass, degradation rate, and microstructural transformation. In Fig. 3a and c, the volume and shape of the 2% CMCTS/3% oHA hydrogel significantly decreased on day 1 compared to day 0, while there were no remarkable changes for the 4% CMCTS/3% oHA hydrogel until day 3. Fig. 3b of degradation kinetics revealed that the 2% CMCTS/3% oHA condition exhibited a rapid degradation rate and almost completely degraded by day 3. In contrast, the 4% CMCTS/3% oHA hydrogel degraded moderately by day 14. As a result, we determined that increased crosslinking intensity could slow down the degradation rate [46]. The internal microstructure of the 4% CMCTS/3% oHA hydrogel was confirmed throughout the entire degradation process, as seen in Fig. 3d and e. Following day 1 of immersion in PBS, the hydrogel swelled significantly, and the average pore size became much larger (93.32 µm) than that of the hydrogels produced on day 0 (60.92 µm), providing a superior three-dimensional extracellular matrix for loading AgSD and DPSCs. After day 2, although the hydrogel maintained a porous structure, the average pore size continued to decrease until the degradation test was completed. Additionally, *in vitro* degradation tests of CMCTS/oHA hydrogels were also conducted in different media (Fig. S4a and b). The results indicated that CMCTS/oHA hydrogels could be completely degraded within 10 days, trends were similar with composite degradation results. Notably, the initial swelling trend of the hydrogels in the FBS and medium groups was significantly lower than in the PBS group. Among the groups, the 10% FBS group exhibited the fastest degradation behavior (Fig. S4b). It is well known that hydrogel degradation can be influenced by various *in vitro* medium conditions, including different enzymatic presence and ionic strength [47, 48]. Specifically, this may be attributed to the fact that the addition of FBS altered the internal structure of the hydrogels. For the comparison of degradation speed between CMCTS/oHA composites and hydrogels, CMCTS/oHA hydrogels showed slightly rapid degradation rate. Based on these results, 4% CMCTS/3% oHA hydrogels were chosen for *in vitro* and *in vivo* trials.

**Fig. 3.**
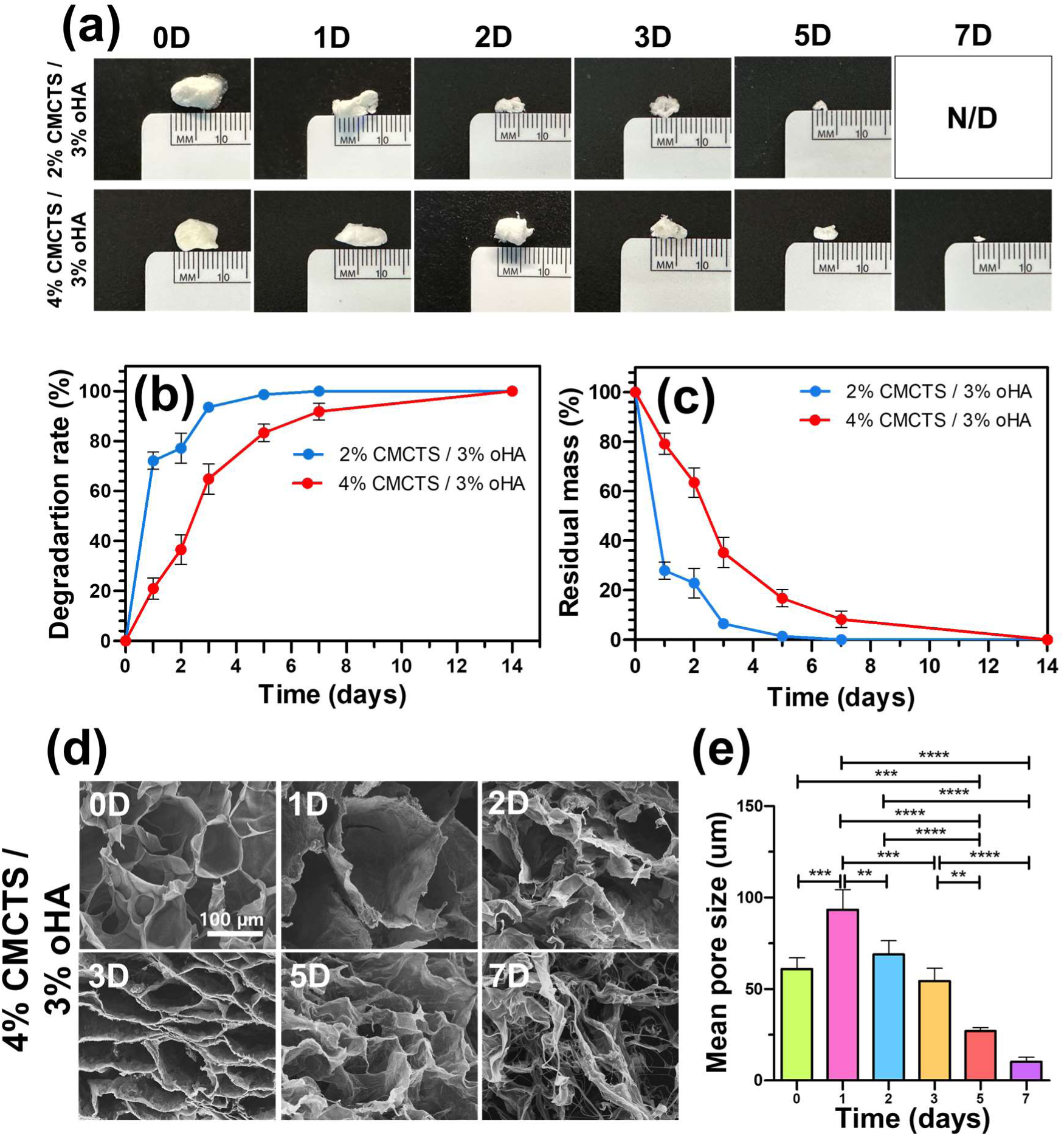
Degradation kinetics of 2% CMCTS / 3% oHA and 4% CMCTS / 3% oHA hydrogels. Optical images of freeze-dried residual hydrogels over time after degradation (a), measurement of degradation rate (b), and residual mass (c). The degradation rate of 2% CMCTS / 3% oHA hydrogels was faster than that of 4% CMCTS / 3% oHA hydrogels. SEM images of freeze-dried 4% CMCTS / 3% oHA hydrogels at 0, 1, 2, 3, 5, and 7 days (d) and average pore sizes (e). Pore sizes gradually decreased over time. Values of *p\** < 0.05, *p\*\** < 0.01, *p\*\*\** < 0.001, and *p\*\*\*\** < 0.0001 were considered statistically significant.

To verify the biocompatibility, high-density DPSCs-laden CMCTS/oHA and CMCTS/oHA + 0.5% AgSD hydrogels were cultured for 7 days to confirm biocompatibility and intracellular behavior. In Fig. 4a, the CMCTS/oHA group was placed in a cell culture plate with medium, and individual cells were entrapped within the hydrogel matrix at 0D. Interestingly, the hydrogel gradually disappeared over time, and high-density cells organized together, forming a 3D tissue resembling a 3D spheroid. Conversely, the CMCTS/oHA + 0.5% AgSD group did not show a similar tendency to form a 3D tissue, with cells dispersed on the cell culture plate. This result suggests that the self-degradation of the CMCTS/oHA hydrogel and cell-to-cell adhesion in the CMCTS/oHA group facilitated 3D cell aggregation, while the CMCTS/oHA + 0.5% AgSD group could not support 3D cell aggregation, likely due to excessive AgSD causing cell death [28]. Consequently, Ag-based materials exhibit cytotoxicity, necessitating a suitable balance in biomaterials. More detailedly, cell viability was confirmed by CCK-8 assay (Fig. 4b). Over 7 days of culture, DPSCs dramatically increased from day 3 to day 7, whereas DPSCs continuously decreased in the CMCTS/oHA + 0.5% AgSD group. This result indicated that 0.5% AgSD exhibited cytotoxicity against DPSCs during culture. On day 7, live and dead DPSCs were visualized (Fig. 4c). The 3D DPSCs aggregate from the CMCTS/oHA group expressed strong green (live) and no red (dead). However, most DPSCs in the CMCTS/oHA + 0.5% AgSD group expressed red (dead) rather than green (live), indicating significant cell death. This result corresponded with microscopic visualization (Fig. 4a) and the CCK-9 assay (Fig. 4b). It is caused by AgSD releases Ag ions within the hydrogel, which exhibit cytotoxic properties. High concentrations of Ag ions can disrupt cell membranes, leading to apoptosis or necrosis [49]. Additionally, Ag ions can induce the production of reactive oxygen species in cells, triggering oxidative stress responses that damage intracellular proteins, lipids, and DNA [40]. Thus, high concentrations of Ag ions reduce cell viability, thereby affecting cell proliferation and aggregation. We verified that 3D cell aggregates were produced from the CMCTS/oHA hydrogel. To check the matrix of the 3D aggregate, histological analysis was performed (Fig. 4d). As expected, individual cells were encapsulated within CMCTS/oHA at 0D. From day 1, DPSCs condensed, finally forming a 3D spheroidal structure on day 7, clearly revealing strong cell-to-cell condensation. As mentioned above, this was facilitated by the self-degradation of CMCTS/oHA, allowing DPSCs to interact with each other. In this study, we emphasize the encapsulation of high-density DPSCs in the CMCTS/oHA hydrogel, and spontaneously formulation of 3D tissue. This phenomenon can be explained by spontaneous and rapid degradation of the hydrogel’s characteristic. In fact, various enzymes are often employed to accelerate hydrogel degradation and promote tissue organization. This approach facilitates faster intercellular bonding (e.g., f-actin adherence) and enhances cell-to-cell interactions [50]. When cells come into contact, they can form junctions and create cell clusters [51, 52]. As numerous cells establish junctions, they collectively organize into a structured 3D cell aggregate, commonly referred to as 3D self-assembly.

**Fig. 4.**
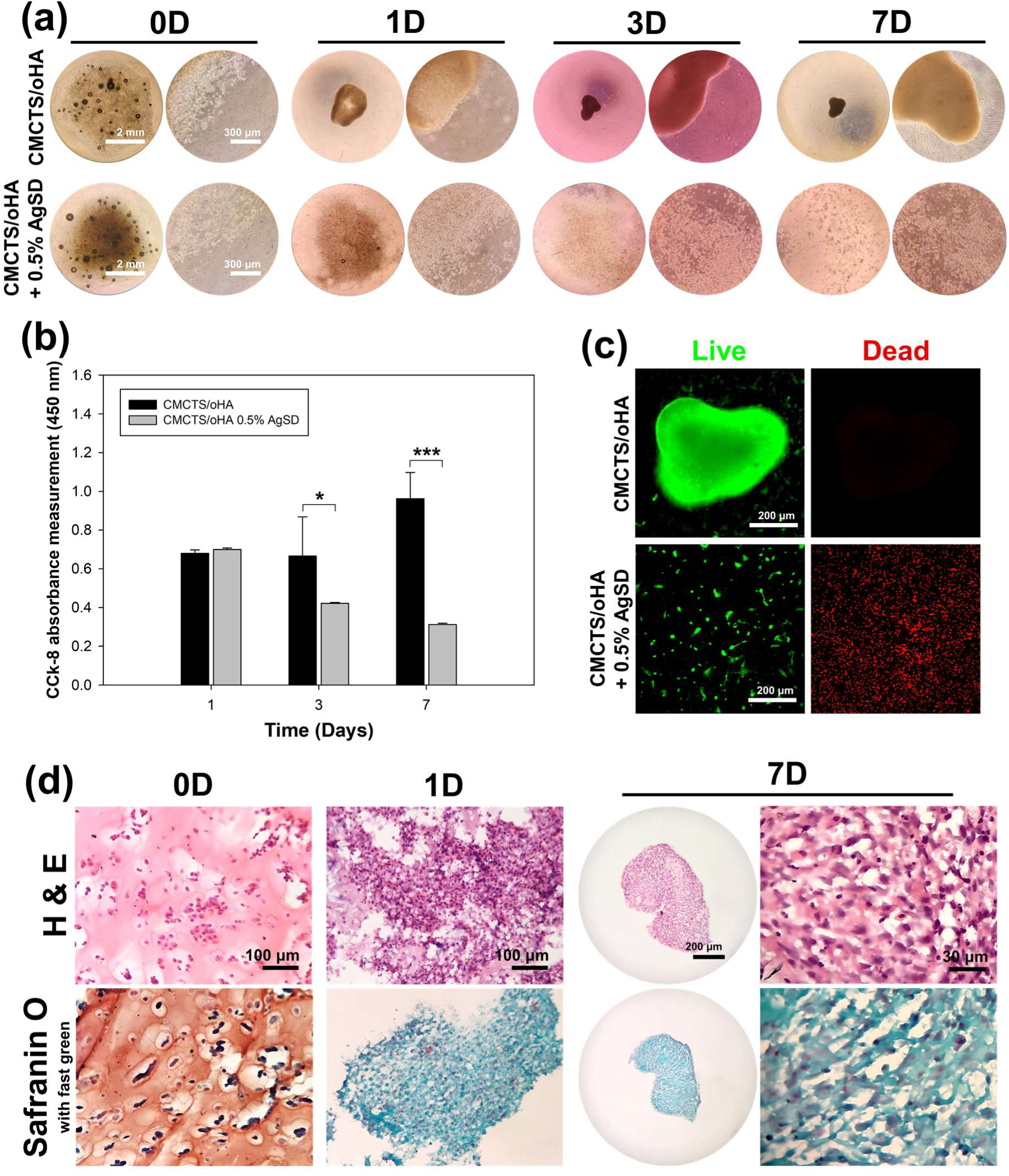
*In vitro* culture of high-density DPSCs-laden CMCTS/oHA and CMCTS/oHA + 0.5% AgSD hydrogels for 7 days. CMCTS/oHA hydrogels facilitated 3D tissue aggregation, whereas CMCTS/oHA + 0.5% AgSD hydrogels did not promote 3D tissue condensation (a). 3D tissue aggregates began to form on day 1 of culture in the CMCTS/oHA group. The cell proliferation rate of CMCTS/oHA and CMCTS/oHA + 0.5% AgSD hydrogels over 7 days (b) and live/dead assay at 7 days of culture (c). Histological analysis of the CMCTS/oHA group at 0, 1, and 7 days of culture using H & E and safranin O with fast green staining (d). 3D tissue condensation was observed on day 1, with strong cell-to-cell communication confirmed by day 7. Safranin O staining was negative from day 1, indicating no residual CMCTS/oHA hydrogels due to self-degradation. Values of *p\** < 0.05 and *p\*\*\** < 0.001 were considered statistically significant.

The remaining CMCTS/oHA with 3D cell interaction was confirmed by safranin O with fast green staining [53]. Compared to 0D, the red color of the acellular hydrogel matrix was not seen from 1 day. Instead, a strong green color was observed, indicating the cell matrix. This result demonstrated that CMCTS/oHA hydrogels degraded by themselves during culture. It is significant that we did not use any enzyme, such as sodium citrate or hyaluronidase, to break down the hydrogel matrix. Generally, hydrogels are manually removed by chemical agents after 3D cell organization within the hydrogel matrix to obtain cell-only condensation [53]. For instance, Torres *et al.* highlighted the use of sodium citrate to degrade their alginate-based bio-printed constructs, releasing 3D spheroids [54]. Calcium ions are removed from the crosslinked chains by citrate cations, destabilizing the alginate constructs. Another study by Wei *et al.* fabricated injectable HA hydrogels with the inoculation of bone marrow mesenchymal stem cell spheroids [55]. Similarly, hydrogels were digested with hyaluronic acid to release 3D cell spheroids after chondrogenic differentiation. Compared to the above examples, the CMCTS/oHA hydrogel system has the great advantage of forming a 3D tissue by self-degradation. As mentioned earlier, the self-degradable CMCTS/oHA hydrogel effectively facilitates the delivery of stem cells and can encapsulate nano/microparticles. Thus, we propose various applications by using CMCTS/oHA hydrogel platform for 3D tissue engineering & regenerative medicine. First, our approach can be induced for facile co-cultured 3D tissue aggregation system by combining multiple cell resources. More specifically, we are able to inject multiple cell resources into the body for promoting tissue formulation. Additionally, our hydrogel can also encapsulate nano/microparticles and deliver them to targeted disease sites, making it suitable for minimally invasive therapies for the complicated conditions such as endodontium or periodontal disease where the operation field is limited in dentistry. Furthermore, it could be clinically utilized for stem cell injections that promote several types of tissue formation such as craniofacial bone cracks. In summary, as it is in the form of an injectable preparation, it can be used as a medical device that does not create surgical scars from incisions. Thus, we believe that our approach has been proven to be highly convenient and enables the facile production of 3D cellular constructs for rapid tissue regeneration.

Further *in vitro* investigations were conducted to evaluate the cytotoxicity of AgSD at concentrations lower than 0.5%. In Fig. 5a, CMCTS/oHA + 0.25% AgSD did not form 3D cell aggregates after 14 days of culture. Instead, cells were stretched into a monolayer on the culture plate (Fig. S5), indicating better cell proliferation compared to the 0.5% AgSD group. The cell proliferation rate in the CMCTS/oHA + 0.1% AgSD group was lower than in the CMCTS/oHA group. However, 3D cell aggregates similar to those in the CMCTS/oHA group were observed after 14 days of culture (Fig. S5). This result suggests that 0.1% AgSD exhibits less cytotoxicity compared to the 0.25% and 0.5% AgSD groups. The CCK-8 assay results showed a gradual increase in cell proliferation for both the CMCTS/oHA and CMCTS/oHA + 0.1% AgSD groups over 7 days. Conversely, the CMCTS/oHA + 0.25% AgSD group displayed poor cell proliferation, indicating that 0.25% AgSD has cytotoxic effects similar to 0.5% AgSD (Fig. 5b). Live/dead assays visualized extensive cell stretching and a high number of live cells in the CMCTS/oHA and CMCTS/oHA + 0.1% AgSD groups, with only a few dead cells (Fig. 5c). The CMCTS/oHA + 0.25% AgSD group exhibited the lowest number of living cells. F-actin staining also demonstrated strong cell stretch and adherence in the CMCTS/oHA and CMCTS/oHA + 0.1% AgSD groups, but few cells in the CMCTS/oHA + 0.25% AgSD group (Fig. 5d). Histological analyses of 3D aggregates in the CMCTS/oHA and CMCTS/oHA + 0.1% AgSD groups revealed similar 3D cell matrices with strong cell-cell connections. In contrast, only a few cells were observed on the culture plate in the CMCTS/oHA + 0.25% AgSD group (Fig. 5e). Based on these results, the optimal AgSD concentration within CMCTS/oHA hydrogels was determined to be 0.1%.

**Fig. 5.**
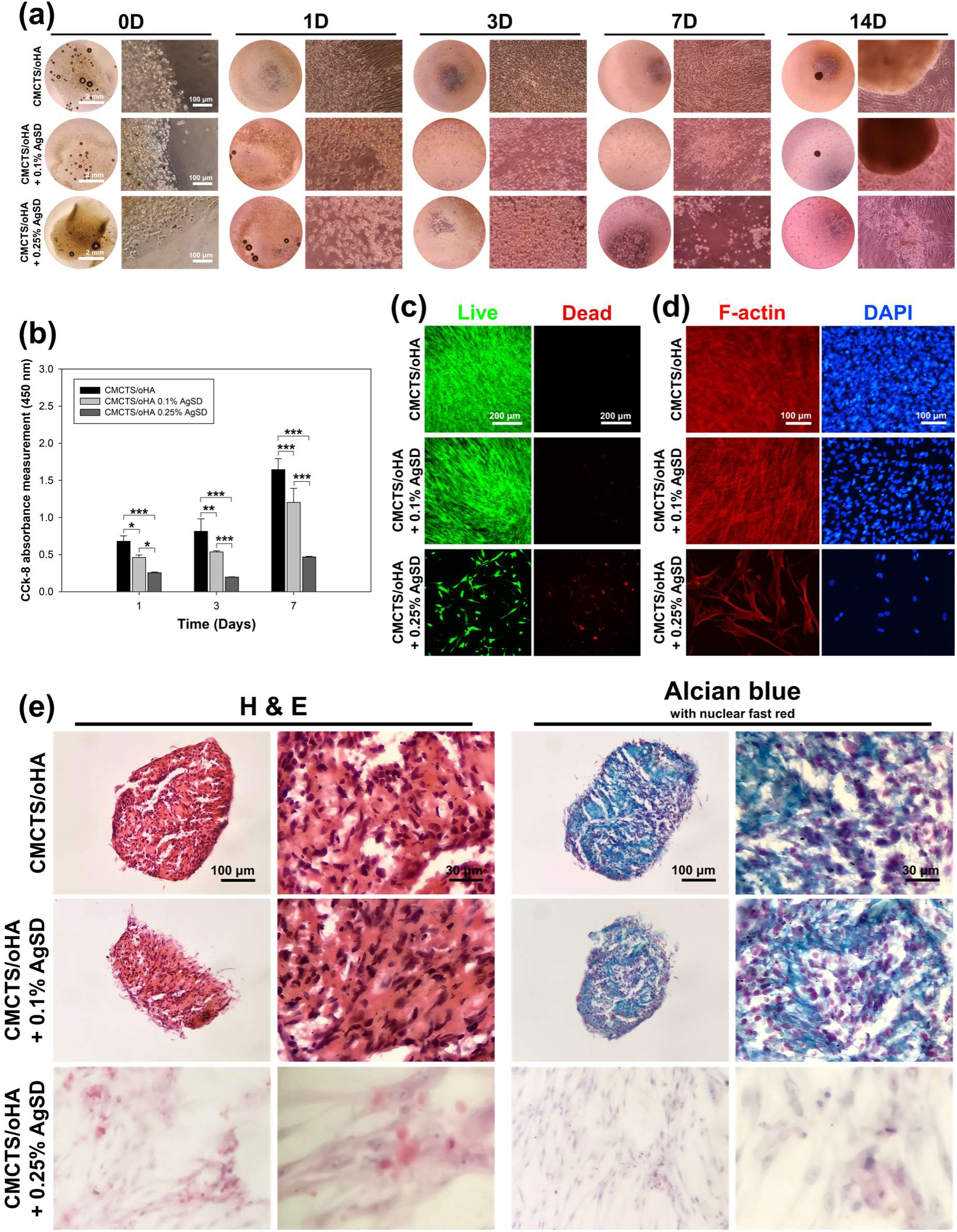
*In vitro* culture of middle-density DPSCs-laden CMCTS/oHA, CMCTS/oHA + 0.1% AgSD, and CMCTS/oHA + 0.25% AgSD hydrogels for 14 days. Cells adhered well to the culture plate and formed 3D tissue aggregates after 14 days of culture in the CMCTS/oHA and CMCTS/oHA + 0.1% AgSD groups (a). DPSCs adhered and proliferated on the culture plate, but 3D tissue condensation did not form in the CMCTS/oHA + 0.25% AgSD hydrogel group. The cell proliferation rate of CMCTS/oHA, CMCTS/oHA + 0.1% AgSD, and CMCTS/oHA + 0.25% AgSD hydrogels over 7 days (b) and live/dead assay (c) with F-actin staining (d) at 7 days of culture. Slight cytotoxicity was observed in CMCTS/oHA + 0.1% AgSD hydrogels compared to CMCTS/oHA hydrogels, but cell proliferation and 3D tissue formation were not impaired. Histological analysis using H & E and alcian blue with nuclear fast red staining revealed strong cell aggregation in the CMCTS/oHA and CMCTS/oHA + 0.1% AgSD groups, but not in the CMCTS/oHA + 0.25% AgSD group. Values of *p\** < 0.05, *p\*\** < 0.01, and *p\*\*\** < 0.001 were considered statistically significant.

In the antibacterial tests, *S. mutans* and *E. faecalis* were employed, given their prevalence in dental caries and infected root canals, respectively [56]. The inhibitory zones for *S. mutans* and *E. faecalis* with different samples are shown in Fig. 6a, with corresponding statistical measurements in Fig. 6b. The negative control (2) and CMCTS/oHA hydrogel (3) exhibited almost no inhibition zones, indicating limited antibacterial effects against *S. mutans* and *E. faecalis*. Among the other samples, the diameter of the antibacterial zones increased in the following order: CMCTS/oHA + 0.1% AgSD hydrogel (4), CMCTS/oHA + 0.25% AgSD hydrogel (5), and TAP (1). For all three samples, the inhibition zones for *S. mutans* were larger than those for *E. faecalis*. These results demonstrate that CMCTS/oHA hydrogels containing AgSD have potent antimicrobial properties against both *S. mutans* and *E. faecalis*, with stronger activity against *S. mutans*. The CMCTS/oHA + 0.25% AgSD hydrogel exhibited superior antibacterial effects compared to the CMCTS/oHA + 0.1% AgSD hydrogel. This indicates that increasing the AgSD content enhances the hydrogel’s antimicrobial activity, confirming that the hydrogel matrix effectively delivers AgSD [57]. Historically, Ag has been recognized as a potent antibacterial agent against *S. mutans* and *E. faecalis* [58, 59].

**Fig. 6.**
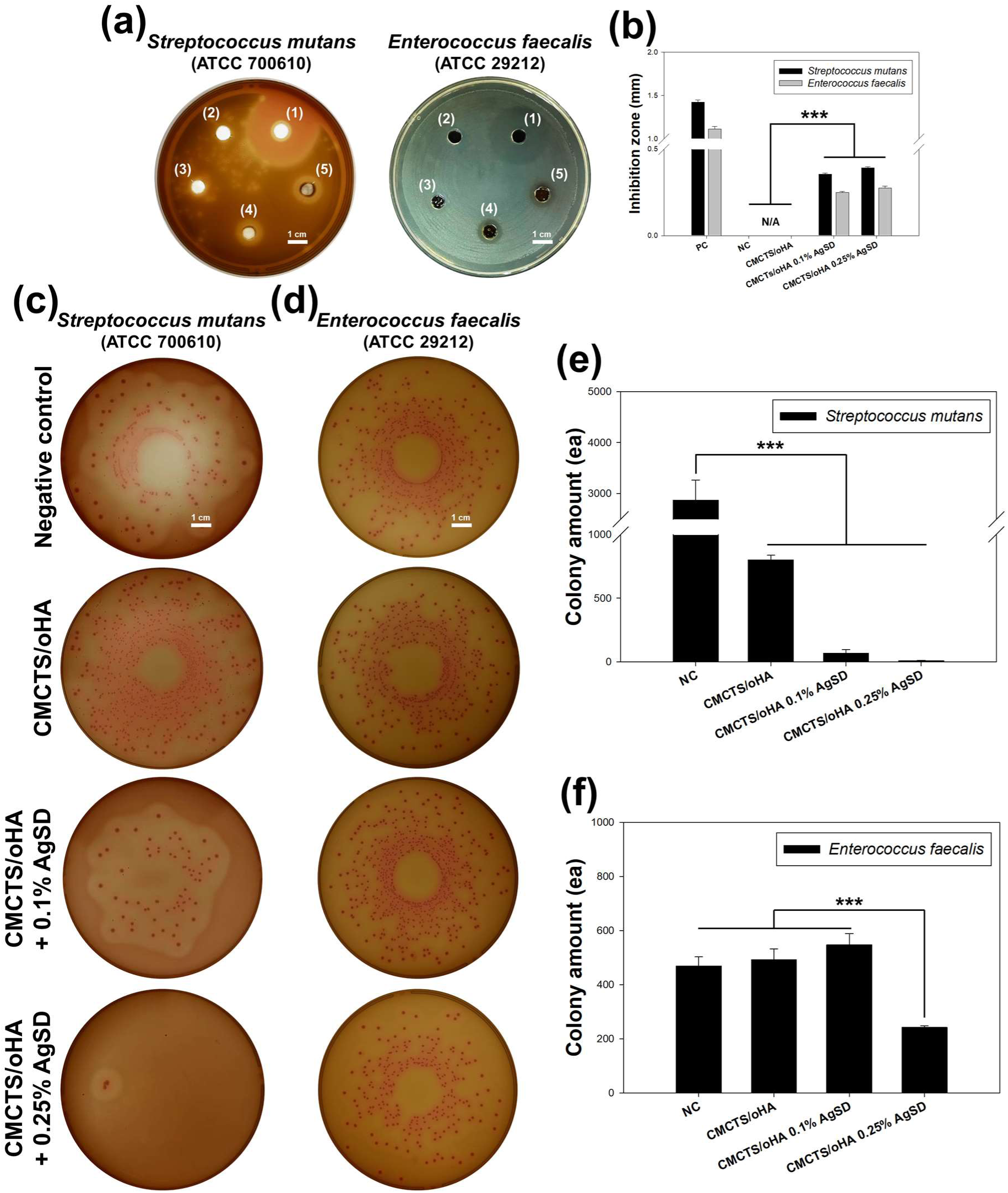
*In vitro* antibacterial test of CMCTS/oHA and CMCTS/oHA/AgSD hydrogels against *S. mutans* and *E. faecalis*. Antibacterial drug as positive control (1), empty as negative control (2), CMCTS/oHA hydrogels (3), CMCTS/oHA + 0.1% AgSD hydrogels (4), and CMCTS/oHA + 0.25% AgSD hydrogels (5) were characterized by inhibition zone test (a) and its quantification result by measuring the diameter of the antibacterial zones (b). The same experimental groups without positive control was also characterized by CFU test (c) and its quantification result (d). The count of *S. mutans* colony was significantly reduced in all experimental groups. However, the colony amount of *E. faecalis* was reduced in CMCTS/oHA + 0.25% AgSD hydrogels group only. Value of *p\*\*\** < 0.001 was considered statistically significant.

The CFU test results are shown in Fig. 6c and e. For *S. mutans*, all CMCTS/oHA conditions significantly reduced bacterial density compared to the negative control. Furthermore, CMCTS/oHA hydrogels containing AgSD reduced bacterial colonies more than the CMCTS/oHA hydrogel alone, indicating that AgSD greatly enhances antibacterial performance against *S. mutans*. This is consistent with the inhibition test findings. For *E. faecalis* (Fig. 6d and f), the CMCTS/oHA hydrogel and CMCTS/oHA + 0.1% AgSD hydrogel groups exhibited slightly increased bacterial colonies compared to the negative control. Significant antibacterial activity was only observed at 0.25% AgSD concentration, which significantly reduced bacterial numbers. This suggests that 0.25% AgSD is necessary for antibacterial effects against *E. faecalis*, as lower concentrations allowed bacterial proliferation. Interestingly, CMCTS/oHA/AgSD hydrogel exhibited better performance against *S. mutans* compared to *E. faecalis.* So far, there have been no systematic comparative studies on the characteristics of the two bacteria. However, we believe their properties are interconnected. The metabolic characteristics of *S. mutans* may make it more sensitive to AgSD. The AgSD exerts its effect by inhibiting the synthesis of folic acid in bacteria, thus *S. mutans* may be more sensitive to this inhibition. On the other hand, *E. faecalis* has more flexible metabolism, with multiple metabolic pathways that allow it to utilize various carbon sources and amino acids as energy. For example, it can use arginine and sorbitol without requiring folic acid, making it less sensitive to AgSD^1,2^. And, *E. faecalis* can withstand extreme environmental conditions such as high salt, high temperature, and low pH, which allows it to survive in various environments. In contrast, *S. mutans* is more suited to acidic environments and conditions rich in sugars. This may explain why AgSD exhibits better antibacterial effects against *S. mutans* ^3,4,5^. Previous research by Sheng *et al.* established a dose-response curve for AgNPs [60]. As Ag+ ion concentration increases, there is a narrow range that stimulates bacterial growth, but beyond this range, antibacterial properties are exhibited. It can be inferred that the Ag+ ions released by CMCTS/oHA + 0.1% AgSD hydrogel were at a concentration range that stimulated bacterial proliferation. Further investigation is required to understand the concentration-dependent antibacterial effects of AgSD and its molecular mechanisms.

In the dentistry of oral bacteria issues, *S. mutans i*s one of the primary pathogens responsible for dental caries. It forms biofilms on the tooth surface and metabolizes carbohydrates to produce organic acids, leading to tooth demineralization and caries formation. When caries are not treated promptly, the bacteria can penetrate the dentin and reach the pulp, causing pulpitis and infection [61]. *E. faecalis* is a common pathogen associated with failed root canal treatments. Clinically, the primary approach to removing infected biofilms within the root canal involves mechanical and chemical cleaning and disinfection. However, due to the complexity of the root canal system and the robust biofilm-forming ability and multidrug resistance of *E. faecalis*, complete eradication is challenging. *E. faecalis* can survive in extreme environments, including high pH and low oxygen conditions, which enhances its survival within the root canal. Its biofilm not only protects the bacteria from antimicrobial agents but also induces persistent inflammation in the pulp and periapical tissues [41]. In these cases, the CMCTS/oHA/AgSD hydrogel, which shows strong antibacterial properties against two bacteria, is convenient for injection and can also carry cells, is considered to have a significant impact on the advancement of dentistry.

To assess the biocompatibility and degradation of CMCTS/oHA/AgSD *in vivo*, hydrogels were injected subcutaneously into mice (Fig. S6). Fig. 7 shows that well-integrated AgSD was found in the 0.25% and 0.5% AgSD groups (red arrows). No adverse tissue reactions such as necrosis or edema were observed in any group, including CMCTS/oHA and CMCTS/oHA/AgSD. H&E staining revealed normal muscle and fat tissue morphology in all groups, with well-preserved connective tissue. Alcian blue with nuclear fast red staining showed similar normal tissue morphology, with no unusual characteristics. All tissues appeared normal and well-integrated, even in the 0.25% and 0.5% AgSD groups, which exhibited cytotoxicity *in vitro*. This suggests that the *in vivo* environment, with its greater complexity and fluid dynamics, may mitigate some cytotoxic effects observed *in vitro*. Therefore, higher AgSD concentrations may be feasible for animal and future clinical applications. However, immune responses and foreign body reactions require further *in vitro* and *in vivo* biological verification. Safranin O with fast green staining indicated complete degradation of CMCTS/oHA hydrogels *in vivo*, confirming their functionality in delivering pharmaceuticals (e.g., AgSD) locally while promoting tissue integration. Major organ tissues exhibited no significant differences or abnormalities compared to the sham group (data not shown), as observed through H&E staining (Fig. 8).

**Fig. 7.**
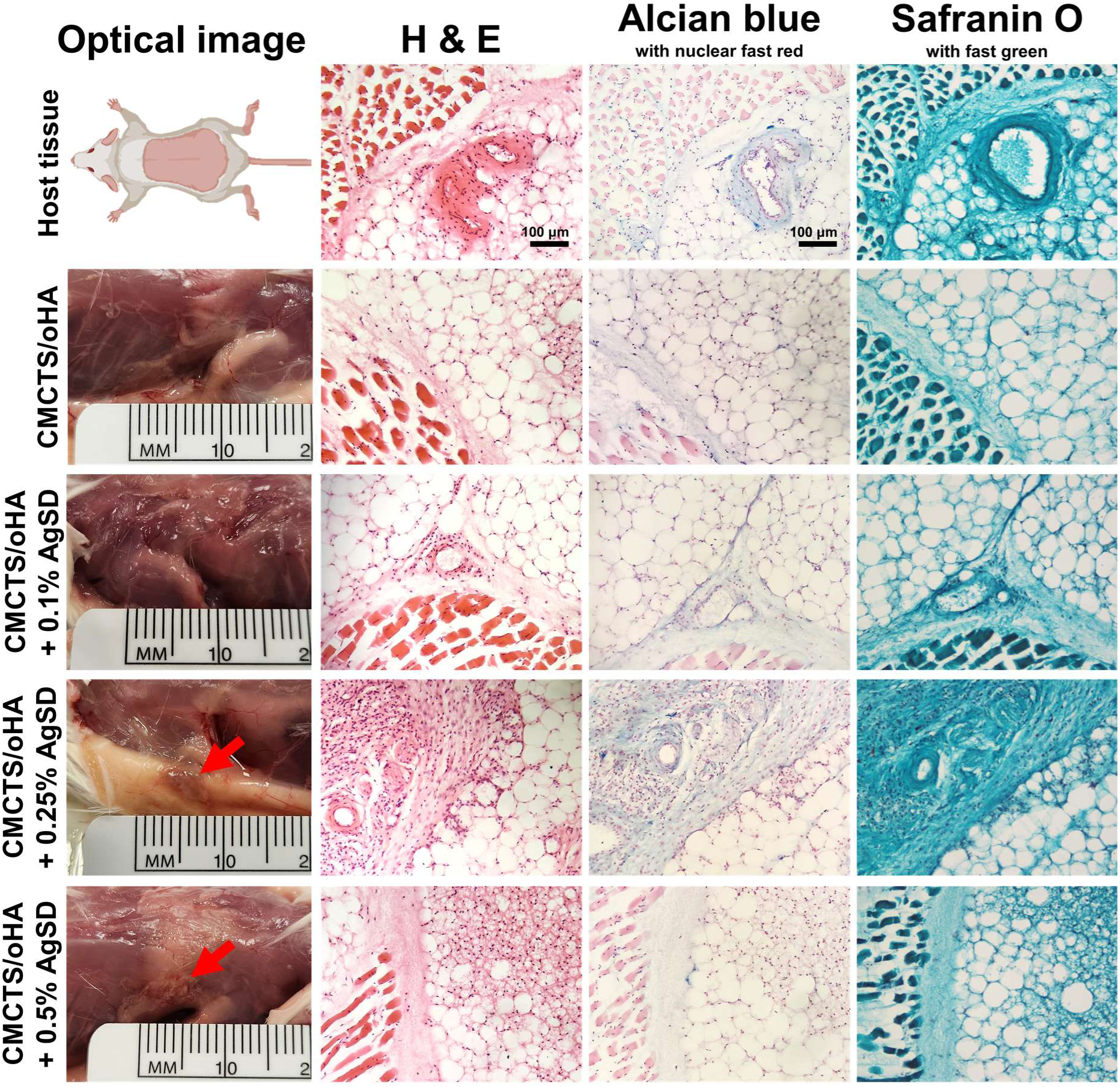
*In situ* injection of CMCTS/oHA, + 0.1% AgSD, + 0.25% AgSD, and + 0.5% AgSD hydrogels into mice subcutaneously and their optical images with histological analysis 2 weeks post-injection. Optical images showed no hydrogel residues or inflammatory pus. Typical mouse subcutaneous tissues, such as adipose, muscle, and connective tissues, were visualized in all staining, with no remaining hydrogels.

**Fig. 8.**
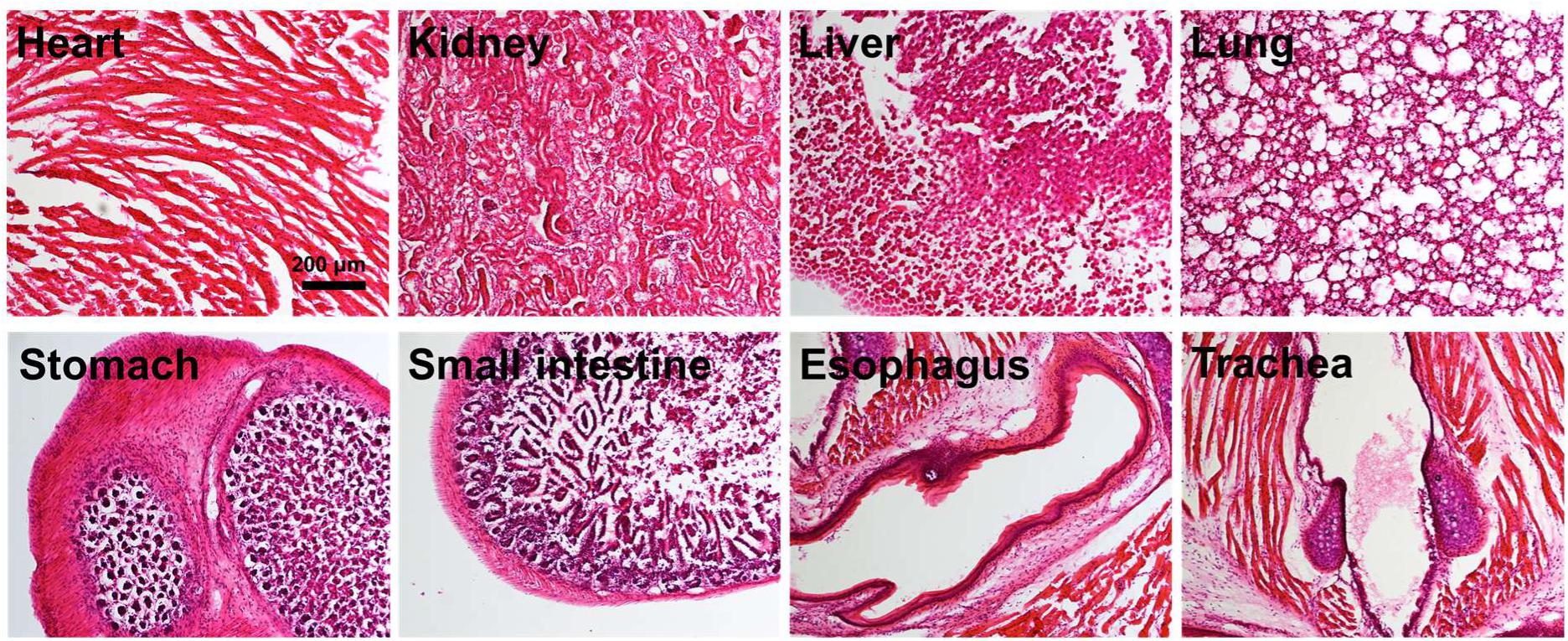
H & E staining of representative organs in the CMCTS/oHA + 0.1% AgSD hydrogel group 2 weeks post-injection. No unusual features were observed, and the tissues appeared similar to those of normal mice organs.

The developed CMCTS/oHA and CMCTS/oHA/AgSD hydrogels demonstrate excellent injectability (Fig. Movies S1 and S2), making them promising candidates for various tissue engineering applications [62]. Their injectability allows for minimally invasive administration, which is advantageous for clinical procedures [21]. For instance, the hydrogel can be precisely delivered to targeted sites, such as periodontal pockets, root canals, or wound beds, using standard syringes. This straightforward injection technology simplifies the clinical application process, reducing the need for complex surgical procedures and enhancing patient comfort [5, 63]. The ease of administration is particularly beneficial in outpatient settings, where quick and effective treatments are essential. As a means of utilizing our system in a hybrid approach, the CMCTS/oHA hydrogel shows promise for the localized delivery of dental stem cells, such as DPSCs and periodontal ligament stem cells, thereby enhancing the regeneration of damaged dental tissues. DPSCs, which are crucial for pulp reconstruction, can be sourced from patients’ own wisdom teeth. When combined with the CMCTS/oHA hydrogel, this system can be effectively used in applications such as dental fillings or restorative materials after pulp capping and pulpotomy, aiming for optimal endodontium regeneration. Maintaining sterility during and after endodontic surgery is vital for prognosis, especially in the microbially rich oral environment. In this context, incorporating antibacterial agents like AgSD helps mitigate infection risks at postoperative sites, promoting faster healing and better pulp recovery outcomes. Moreover, in the context of wound healing—particularly for craniofacial injuries— the hydrogel’s ability to maintain a moist microenvironment while promoting cell aggregation can significantly accelerate the repair of both soft and hard tissue damage. Its injectable formulation allows for minimally invasive applications, which is advantageous for managing chronic wounds or burns, where traditional treatments may be less effective.

Additionally, long-term biocompatibility is essential for therapeutic applications, particularly in dentistry and craniofacial wound healing. Typically, the hydrogel remains in contact with tissues for extended periods. We recognize the need for thorough *in vivo* studies to evaluate potential inflammatory responses and adverse effects over time. Our investigation shows that the CMCTS/oHA hydrogel degrades quickly, with no residual hydrogels observed *in vivo* (Fig. 7 and S2). Therefore, we propose to address concerns regarding residual hydrogel presence by leveraging its rapid self-degradation properties, which facilitate efficient breakdown in the body and reduce the risk of long-term accumulation for clinical wound healing applications. Scalability related to hydrogel manufacturing for clinical or commercial applications is also a concern. While our formulation shows promise in laboratory settings, translating it into a clinically viable product requires careful optimization of production techniques to ensure consistent quality and efficacy. Our approach, which utilizes minimal chemical modifications, enhances reproducibility and aligns with regulatory requirements. Through our straightforward production methods, we have established a robust framework for transitioning from laboratory to clinical settings, enabling the widespread use of our hydrogel in various therapeutic applications. By incorporating these considerations, we aim to provide a balanced perspective that acknowledges both the potential benefits and the challenges associated with the clinical use of our CMCTS/oHA hydrogel system.

The *in vivo* studies confirmed that CMCTS/oHA hydrogels degrade completely, which is a critical feature for tissue engineering applications [64]. The degradation of the hydrogel ensures that it does not require surgical removal after fulfilling its therapeutic purpose. This property is highly desirable in regenerative medicine, where temporary scaffolds are needed to support tissue growth and then naturally degrade to allow for new tissue formation [65]. The ability of CMCTS/oHA hydrogels to form 3D cell aggregates and support cell proliferation makes them suitable for 3D tissue engineering. These hydrogels can be used to create 3D tissue constructs for repairing or replacing damaged tissues and organs. For instance, the hydrogel can be used to engineer cartilage, bone, or skin tissues, providing a scaffold that promotes cell growth and tissue regeneration [66]. The incorporation of AgSD provides antimicrobial properties, which is particularly useful in preventing infections during the healing process. Thus, the CMCTS/oHA hydrogel, with its excellent injectability, controlled degradation, and ability to support 3D tissue engineering, represents a significant advancement in bioengineering. Its potential applications in clinical practice are vast, offering improved therapeutic outcomes and patient experiences across a range of medical disciplines in near future.

## 4. Conclusion

In summary, this study presents the AgSD incorporated CMCTS/oHA hydrogel as a promising biomaterial for diverse tissue engineering applications. The hydrogel’s excellent injectability allowed for minimally invasive administration, enhancing an applicability and hand-comfort. *In vitro* assessments identified 0.1% AgSD as the optimal concentration for balancing cell proliferation and antimicrobial efficacy. Antibacterial tests demonstrated significant inhibitory effects against *S. mutans* and *E. faecalis*, with increased AgSD concentrations enhancing antimicrobial activity. *In vivo* evaluations confirmed the CMCTS/oHA hydrogel’s biocompatibility and complete degradation, ensuring no adverse tissue reactions and supporting tissue integration. These findings underscore the hydrogel’s suitability for 3D tissue engineering, enabling the creation of 3D cell condensations. The CMCTS/oHA hydrogel’s showed self-crosslinking without additional crosslinker, self-degradation, antimicrobial activity, and ability to support 3D tissue formation as a valuable tool in regenerative medicine. Future studies would focus on further optimizing the hydrogel’s properties and exploring its efficacy for clinical settings, paving the way for its integration into routine medical practice and enhancing patient care.

## Acknowledgement

This work was supported by the Basic Science Research Program through the National Research Foundation of Korea (NRF) funded by the Ministry of Science and ICT (RS-2024-00338610). This work was supported by the National R&D Program through the National Research Foundation of Korea (NRF) funded by the Ministry of Science and ICT (RS-2024-00405574).

