## Supplementary for "Crosslinker-free *in situ* hydrogel induces self-aggregation of human dental pulp stem cells with enhanced antibacterial activity"

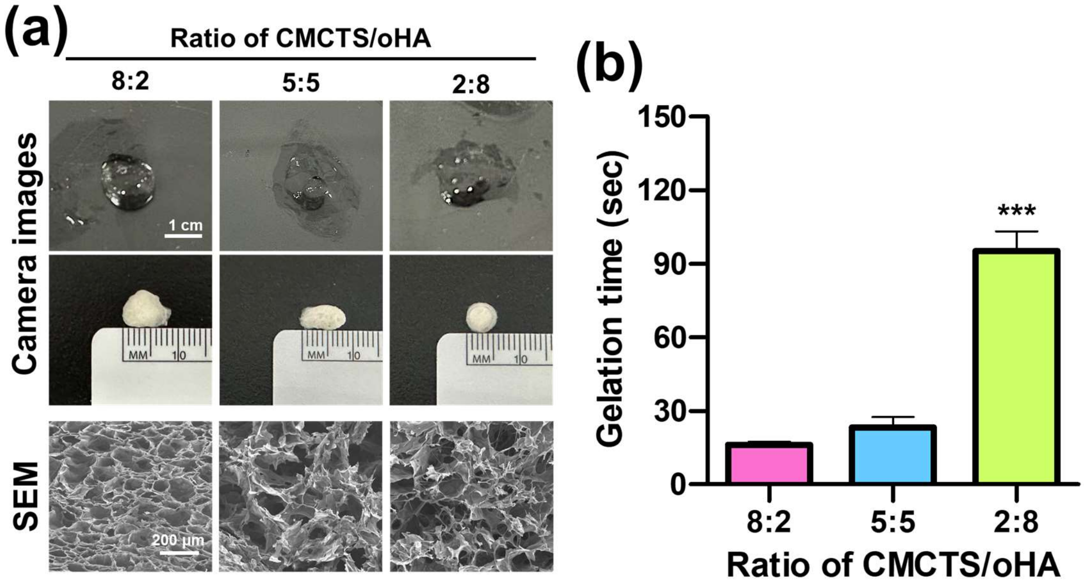


**Fig. S1.** Preparation of 4% CMCTS / 3% oHA hydrogels at different ratios of 8:2, 5:5, and 2:8. Hydrogels were prepared in clear plates and showed different fabrication phenomena (a). The SEM image demonstrated that the 8:2 mixing condition had the most uniform pore structure. The gelation time measurement results indicated that the 8:2 condition formed a complete hydrogel (b). Values of $p^{***} < 0.001$ were considered statistically significant against 8:2 and 5:5.

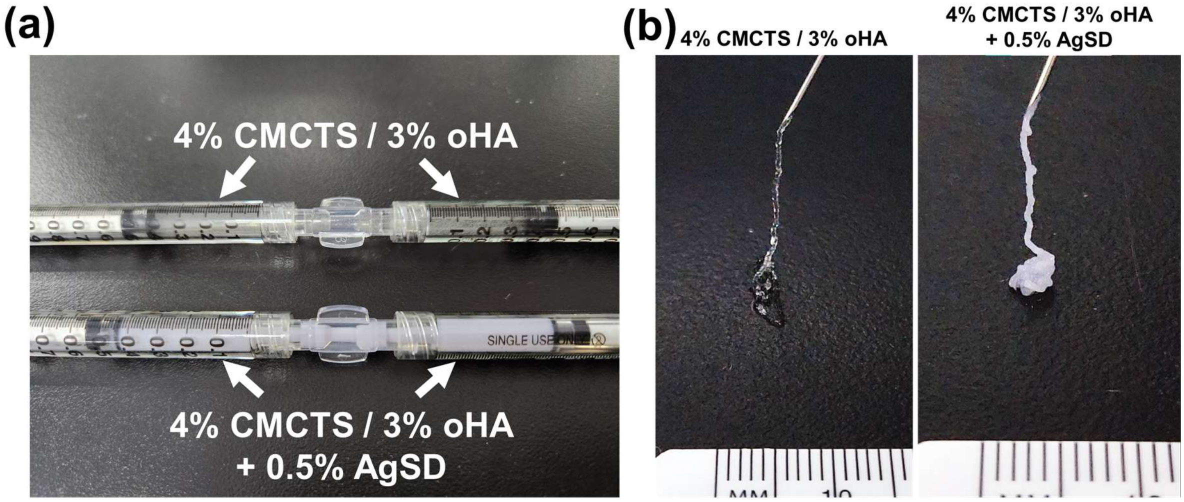


**Fig. S2.** Preparation of 4% CMCTS / 3% oHA and 4% CMCTS / 3% oHA + 0.5% AgSD hydrogels was achieved. The tunnel mixing system enabled gelation without the need for a crosslinking agent, occurring at room temperature. Both hydrogel formulations were well-conjugated (a). When administering the hydrogels using a 27-gauge syringe, the injection was smoothly executed while preserving the desired viscosity (b). The inclusion of AgSD did not adversely impact the injectability of the hydrogels.

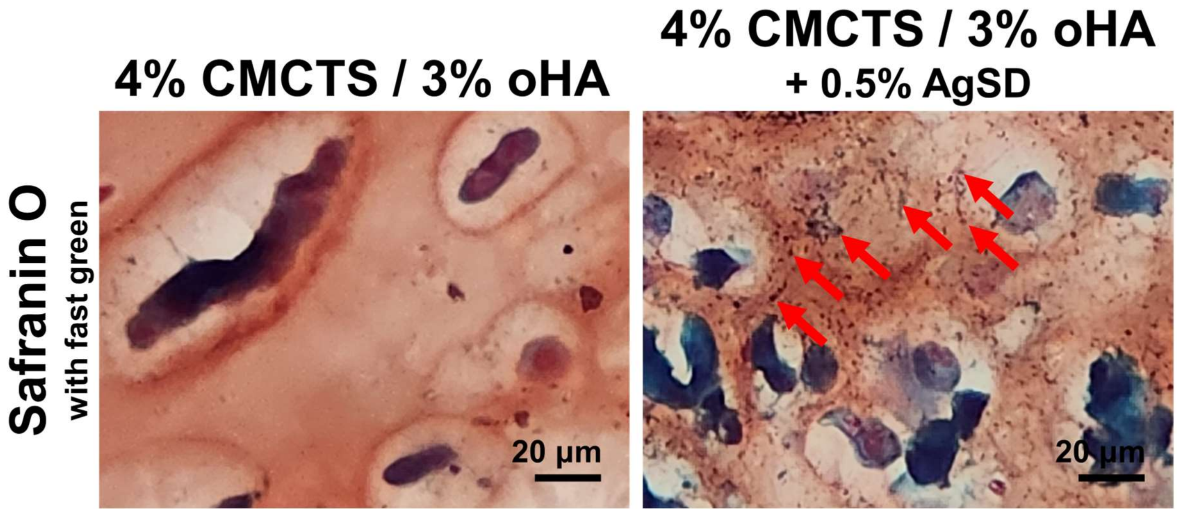


**Fig. S3.** Safranin O staining of the 4% CMCTS / 3% oHA and 4% CMCTS / 3% oHA + 0.5% AgSD hydrogels at day 0. The hydrogels were immediately fixed and subjected to the staining procedure upon conjugation. The cells were stained green through the application of fast green, while the CMCTS/oHA matrix exhibited a vibrant red coloration. Within the 4% CMCTS / 3% oHA + 0.5% AgSD hydrogels, discrete particulate matter consistent with AgSD was observed, which was well-immobilized within the hydrogel network. The encapsulated cells were also firmly embedded throughout the hydrogel structure.

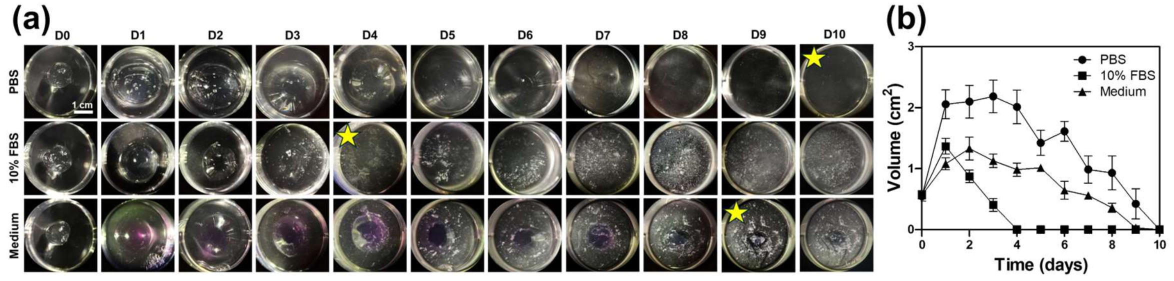


**Fig. S4.** Degradation test of 4% CMCTS / 3% oHA under 1x PBS, 10% FBS, and completed cell growth medium for 10 days (a). Degradation trends were measured using the remaining hydrogel volume (b). The degradation of the CMCTS/oHA hydrogel varied depending on the environment, and the point of complete degradation was indicated by a yellow star.

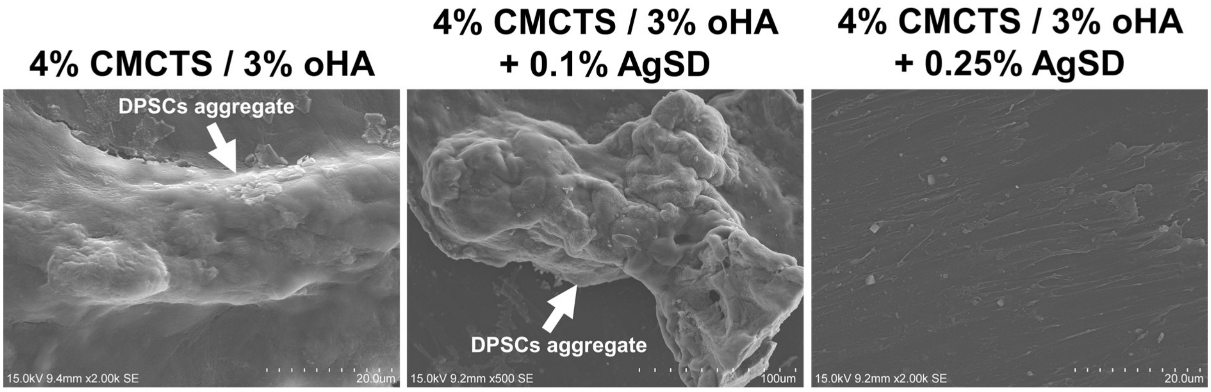


**Fig. S5.** SEM imaging revealed the cellular aggregation patterns. Within the CMCTS/oHA and CMCTS/oHA + 0.1% AgSD groups, robust cellular aggregation was observed. In contrast, the 0.25% AgSD group exhibited a limited number of sparsely distributed cells adhered to the culture plate surface.

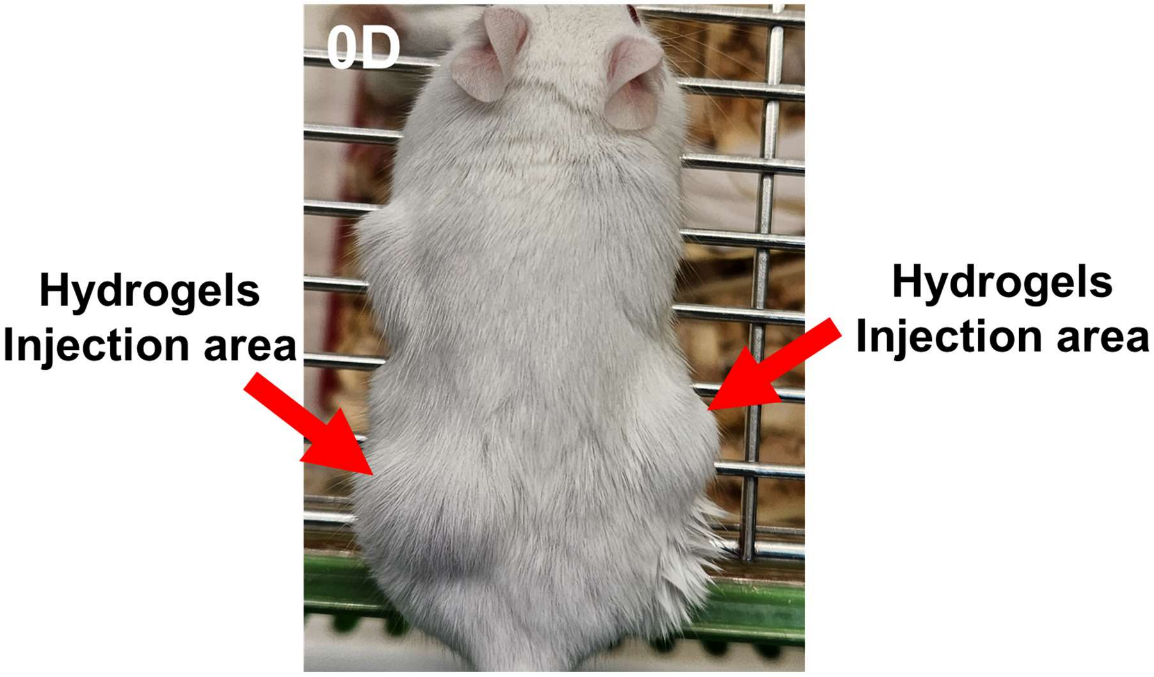


**Fig. S6** Immediately following hydrogel administration, the appearance of the mice is shown. The hydrogel was injected subcutaneously on the dorsum of the mice without any prior incision. The injected hydrogel maintained its volumetric integrity.
